## Supplemental Figures for "Laminin γ1-dependent basement membranes are instrumental to ensure proper olfactory placode shape, position and boundary with the brain, as well as olfactory axon development"

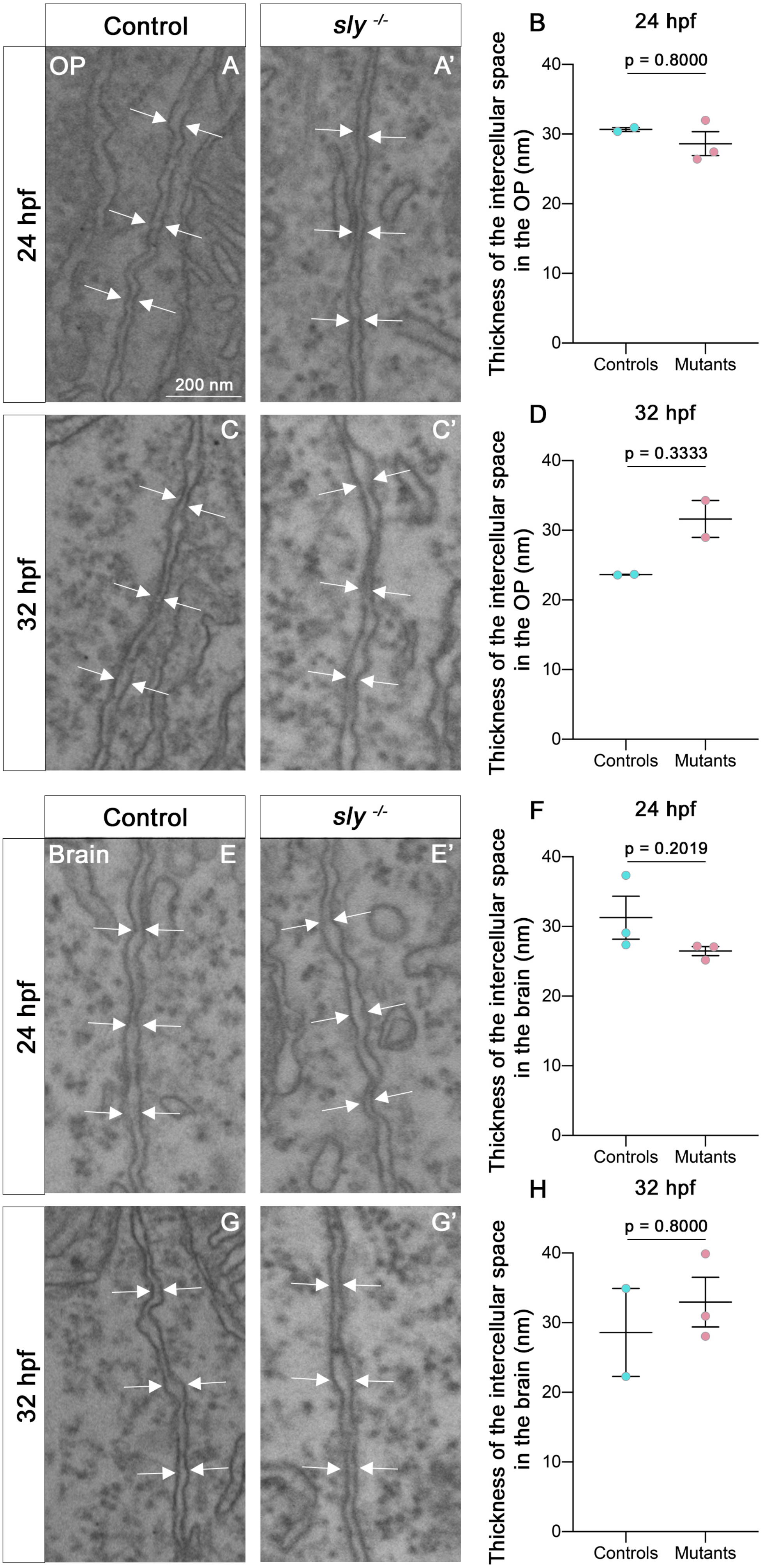

Control
sly -/-
OP
A
A'
24 hpf
200 nm
C
C'
32 hpf
Brain
E
E'
24 hpf
G
G'
32 hpf
B
24 hpf
p = 0.8000
Thickness of the intercellular space in the OP (nm)
Controls
Mutants
D
32 hpf
p = 0.3333
Thickness of the intercellular space in the OP (nm)
Controls
Mutants
F
24 hpf
p = 0.2019
Thickness of the intercellular space in the brain (nm)
Controls
Mutants
H
32 hpf
p = 0.8000
Thickness of the intercellular space in the brain (nm)
Controls
Mutants


Figure S1

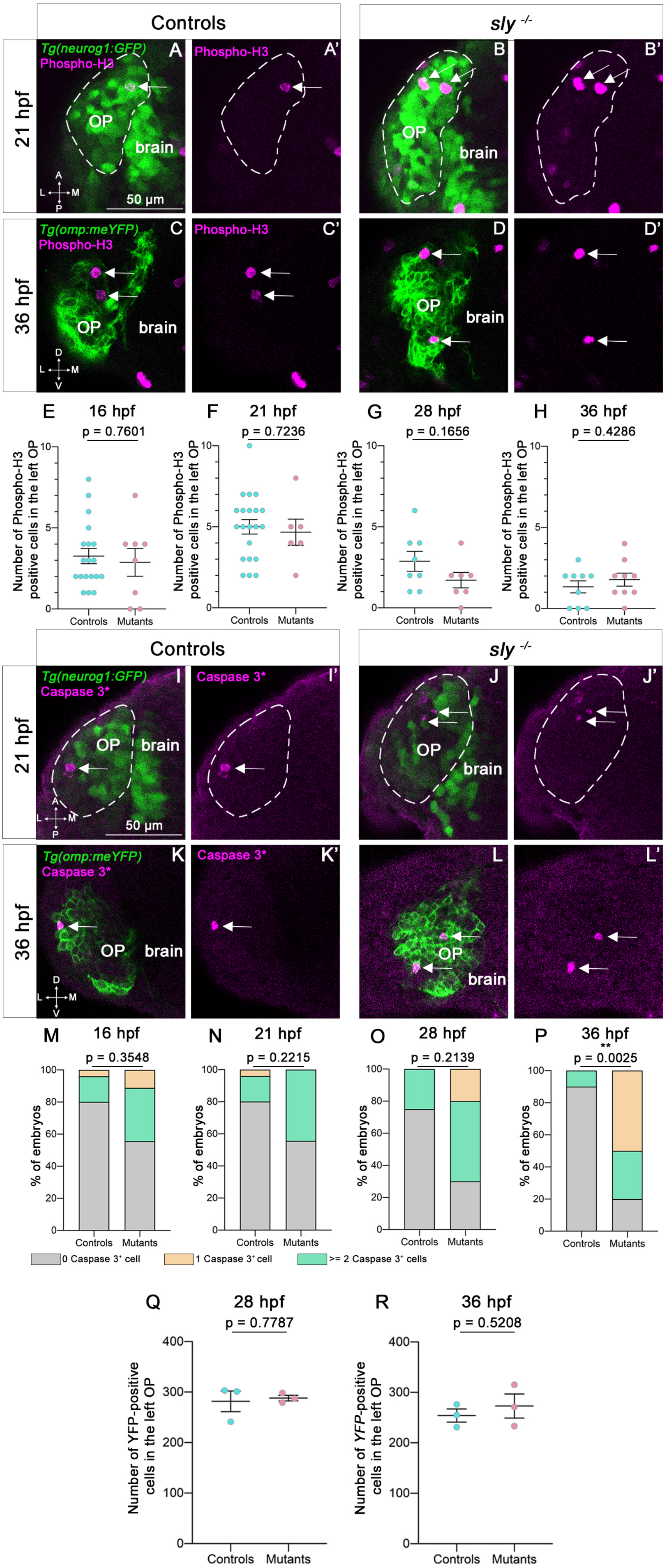
Controls
sly -/-
21 hpf
36 hpf
Tg(neurog1:GFP)
Phospho-H3
A
A'
B
B'
OP
brain
50 µm
Tg(omp:meYFP)
C
C'
D
D'
E 16 hpf p = 0.7601
F 21 hpf p = 0.7236
G 28 hpf p = 0.1656
H 36 hpf p = 0.4286
Number of Phospho-H3 positive cells in the left OP
Controls
Mutants
Caspase 3*
I
I'
J
J'
K
K'
L
L'
M 16 hpf p = 0.3548
N 21 hpf p = 0.2215
O 28 hpf p = 0.2139
P 36 hpf ** p = 0.0025
% of embryos
0 Caspase 3⁺ cell
1 Caspase 3⁺ cell
>= 2 Caspase 3⁺ cells
Q 28 hpf p = 0.7787
R 36 hpf p = 0.5208
Number of YFP-positive cells in the left OP

Figure S2

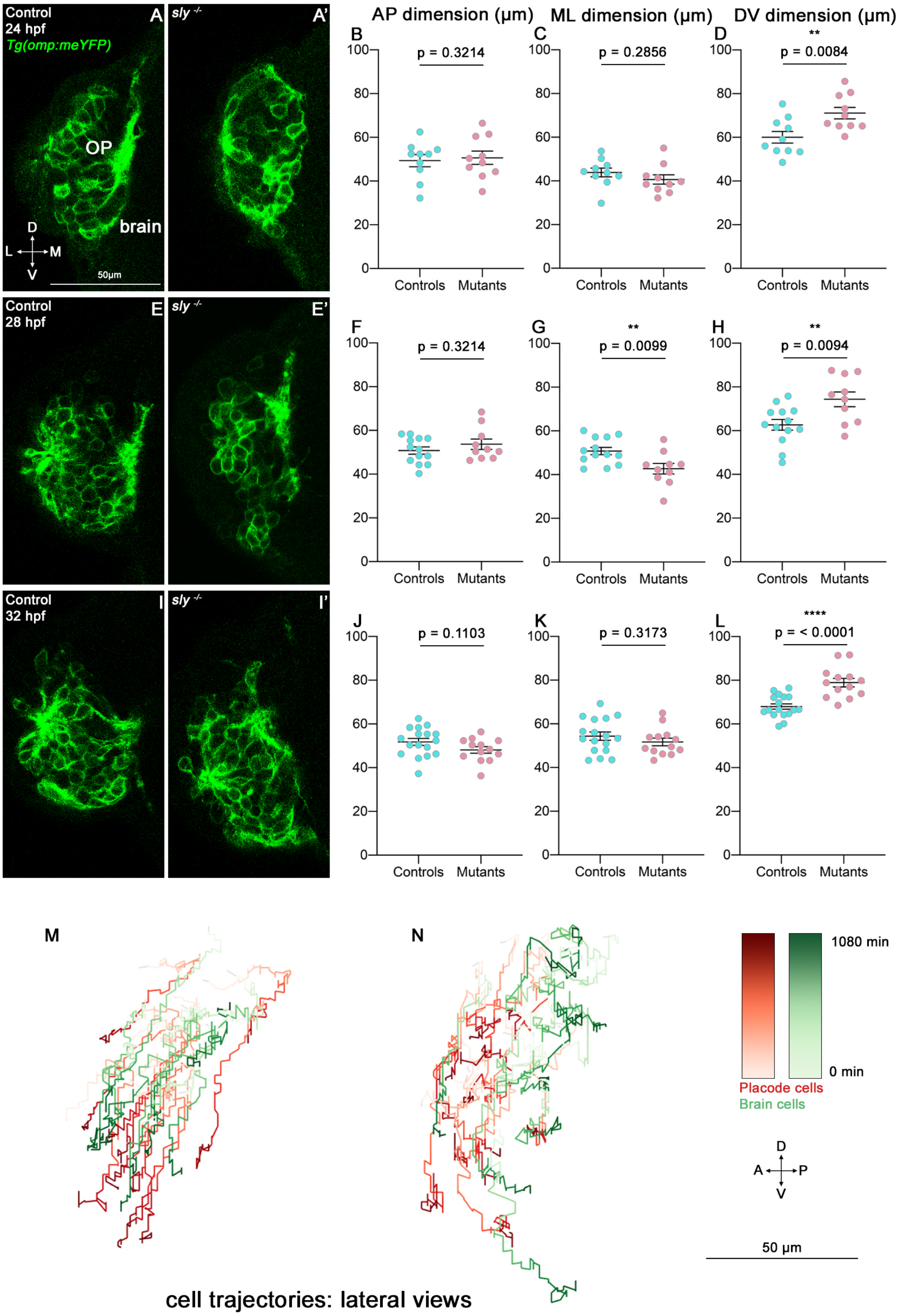

Control
24 hpf
Tg(omp:meYFP)
A
OP
brain
D
L
M
V
50μm
sly -/-
A'
AP dimension (μm)
ML dimension (μm)
DV dimension (μm)
B
p = 0.3214
C
p = 0.2856
D
**
p = 0.0084
Controls
Mutants
Control
28 hpf
E
sly -/-
E'
F
p = 0.3214
G
**
p = 0.0099
H
**
p = 0.0094
Control
32 hpf
I
sly -/-
I'
J
p = 0.1103
K
p = 0.3173
L
****
p = < 0.0001
M
N
1080 min
0 min
Placode cells
Brain cells
D
A
P
V
50 μm
cell trajectories: lateral views


### Figure S3

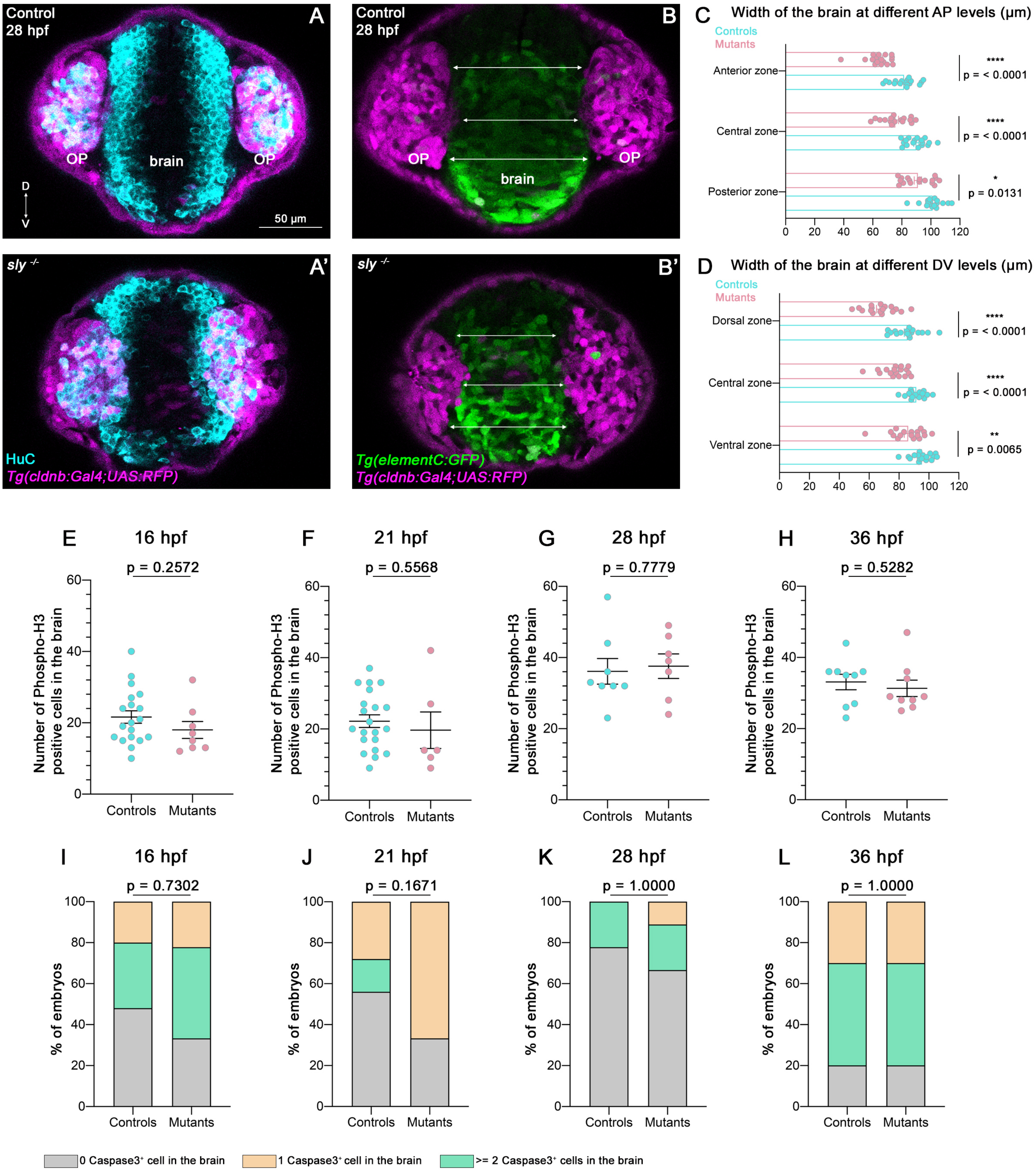
Control
28 hpf
A
OP
brain
OP
D
V
50 µm
Control
28 hpf
B
OP
brain
OP
C
Width of the brain at different AP levels (µm)
Controls
Mutants
Anterior zone
Central zone
Posterior zone
****
p = < 0.0001
****
p = < 0.0001
*
p = 0.0131
0 20 40 60 80 100 120
sly -/-
A'
HuC
Tg(cldnb:Gal4;UAS:RFP)
sly -/-
B'
Tg(elementC:GFP)
Tg(cldnb:Gal4;UAS:RFP)
D
Width of the brain at different DV levels (µm)
Controls
Mutants
Dorsal zone
Central zone
Ventral zone
****
p = < 0.0001
****
p = < 0.0001
**
p = 0.0065
0 20 40 60 80 100 120
E
16 hpf
p = 0.2572
F
21 hpf
p = 0.5568
G
28 hpf
p = 0.7779
H
36 hpf
p = 0.5282
Number of Phospho-H3 positive cells in the brain
0 20 40 60
Controls
Mutants
I
16 hpf
p = 0.7302
J
21 hpf
p = 0.1671
K
28 hpf
p = 1.0000
L
36 hpf
p = 1.0000
% of embryos
0 20 40 60 80 100
Controls
Mutants
0 Caspase3⁺ cell in the brain
1 Caspase3⁺ cell in the brain
>= 2 Caspase3⁺ cells in the brain

### Figure S4

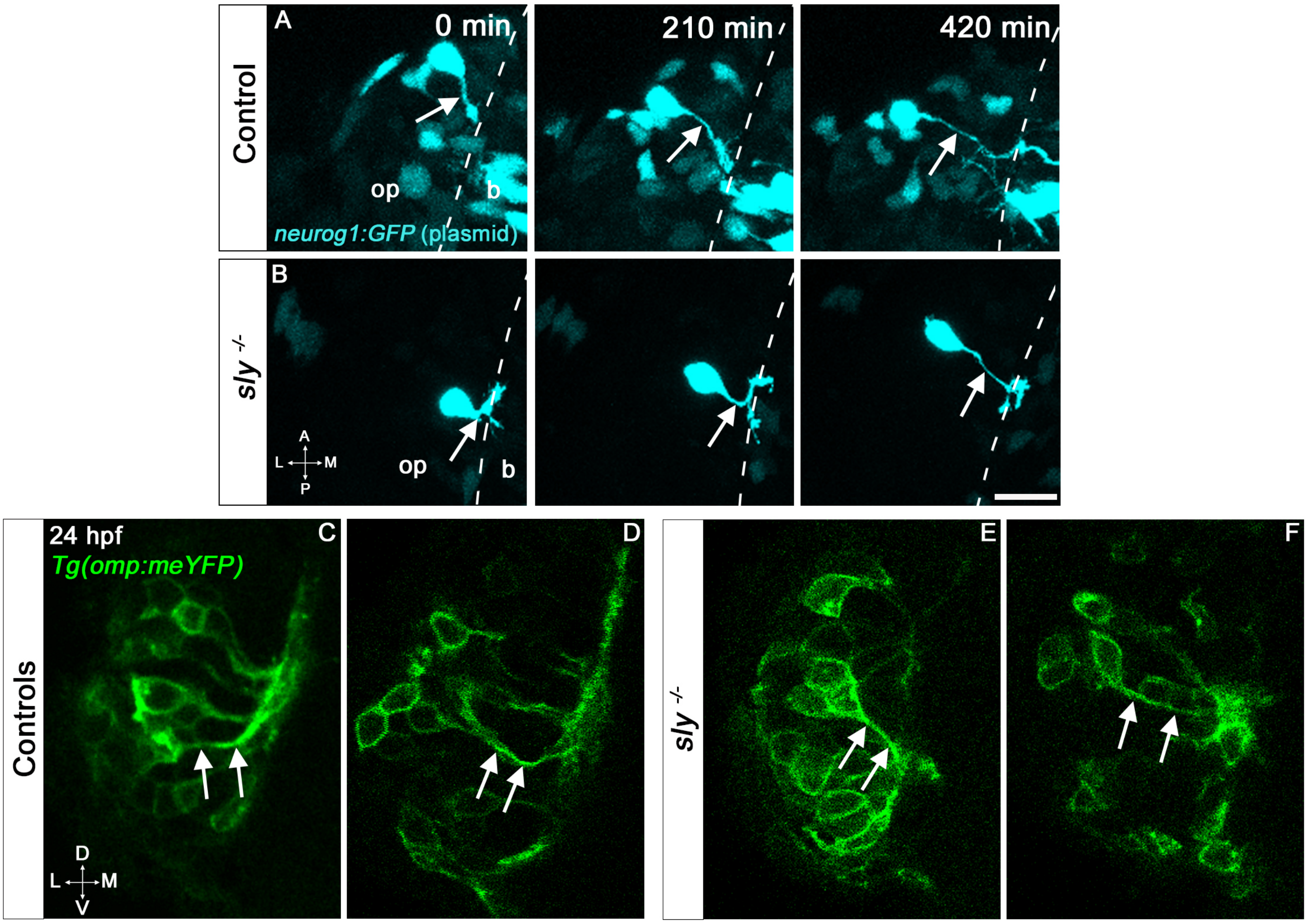


G

| % of embryos with | | A proper axon bundle (dorsal exit point in mutants) | At least one ventral projection | At least one medial projection |
|---|---|---|---|---|
| 24 hpf | Controls (n = 11) | 27 % | 27 % | 9 % |
| | Mutants (n = 10) | 0 % | 50 % | 30 % |
| 28 hpf | Controls (n = 14) | 86 % | 71 % | 0 % |
| | Mutants (n = 11) | 9 % | 73 % | 18 % |
| 32 hpf | Controls (n = 17) | 100 % | 88 % | 0 % |
| | Mutants (n = 13) | 8 % | 92 % | 23 % |
| 36 hpf | Controls (n = 16) | 94 % | 81 % | 6 % |
| | Mutants (n = 10) | 10 % | 90 % | 20 % |

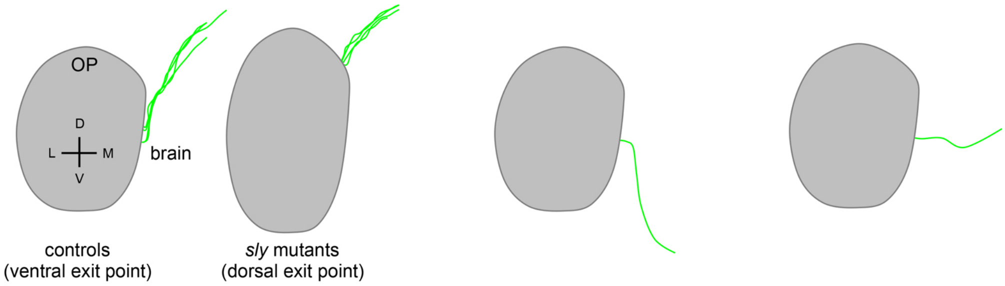


Figure S5

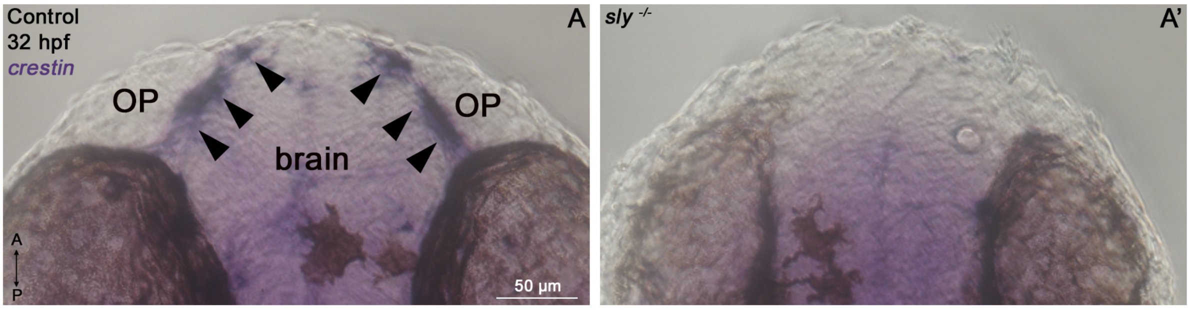


| B Live imaging of NCC migration (mosaïc NCC labelling) | presence of migrating NCC at the **eye/OP interface** | **forebrain/OP interface** | |
|---|---|---|---|
| | | presence of ventral NCC | presence of dorsal NCC |
| **control siblings** | 10/10 placodes | 4/10 placodes<br>In these 4 samples, NCC populated the boundary and stayed there | 8/10 placodes |
| ***sly* mutants** | 7/8 placodes | 2/8 placodes<br>In these 2 samples, NCC migrated at the boundary at early stages but were expelled from it later | 2/8 placodes |

### Figure S6

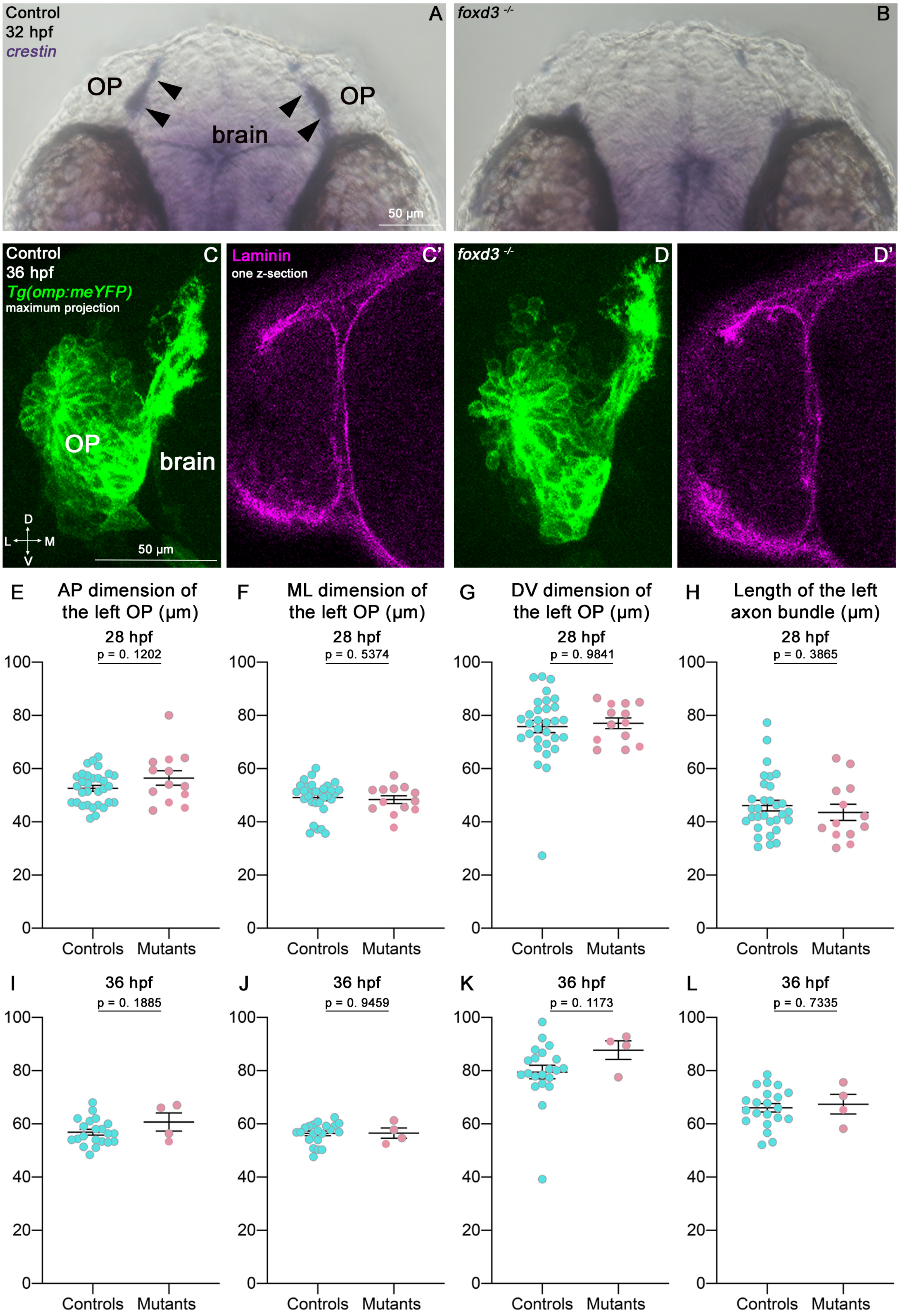
Control
32 hpf
crestin
A
OP
OP
brain
50 µm
foxd3 -/-
B
Control
36 hpf
Tg(omp:meYFP)
maximum projection
C
OP
brain
D
L
M
V
50 µm
Laminin
one z-section
C'
foxd3 -/-
D
D'
E
AP dimension of the left OP (µm)
28 hpf
p = 0. 1202
F
ML dimension of the left OP (µm)
28 hpf
p = 0. 5374
G
DV dimension of the left OP (µm)
28 hpf
p = 0. 9841
H
Length of the left axon bundle (µm)
28 hpf
p = 0. 3865
I
36 hpf
p = 0. 1885
J
36 hpf
p = 0. 9459
K
36 hpf
p = 0. 1173
L
36 hpf
p = 0. 7335
0
20
40
60
80
100
Controls
Mutants

Figure S7
